# Photoacoustic imaging reveals mechanisms of rapid-acting insulin formulations dynamics at the injection site

**DOI:** 10.1101/2022.03.10.483309

**Authors:** Anjul Khadria, Chad D. Paavola, Konstantin Maslov, Francisco A. Valenzuela, Andrea E. Sperry, Amy L. Cox, Rui Cao, Junhui Shi, Patricia L. Brown-Augsburger, Emmanuel Lozano, Ross L. Blankenship, Ranajoy Majumdar, Scott A. Bradley, John M. Beals, Sunday S. Oladipupo, Lihong V. Wang

## Abstract

**Objective:** Ultra-rapid insulin formulations control postprandial hyperglycemia; however, inadequate understanding of injection site absorption mechanisms is limiting further advancement. We used photoacoustic imaging to investigate the injection site dynamics of dye-labeled insulin lispro in the Humalog^®^ and Lyumjev^®^ formulations using the murine ear cutaneous model and correlated it with results from unlabeled insulin lispro in pig subcutaneous injection model.

**Methods:** We employed dual-wavelength optical-resolution photoacoustic microscopy to study the absorption and diffusion of the near-infrared dye-labeled insulin lispro in the Humalog and Lyumjev formulations in mouse ears. We mathematically modeled the experimental data to calculate the absorption rate constants and diffusion coefficients. We studied the pharmacokinetics of the unlabeled insulin lispro in both the Humalog and Lyumjev formulations as well as a formulation lacking both the zinc and phenolic preservative in pigs. The association state of insulin lispro in each of the formulations was characterized using SV-AUC and NMR spectroscopy.

**Results:** Through experiments using murine and swine models, we show that the hexamer dissociation rate of insulin lispro is not the absorption rate-limiting step. We demonstrated that the excipients in the Lyumjev formulation produce local tissue expansion and speed both insulin diffusion and microvascular absorption. We also show that the diffusion of insulin lispro at the injection site drives its initial absorption; however, the rate at which the insulin lispro crosses the blood vessels is its overall absorption rate-limiting step.

**Conclusions:** This study provides insights into injection site dynamics of insulin lispro and the impact of formulation excipients. It also demonstrates photoacoustic microscopy as a promising tool for studying protein therapeutics. The results from this study address critical questions around the subcutaneous behavior of insulin lispro and the formulation excipients, which could be useful to make faster and better controlled insulin formulations in the future.

**Highlights:**

- Hexamer dissociation is not the absorption rate-limiting step for insulin lispro
- Lyumjev excipients enhance insulin microvascular absorption and diffusion
- Vascular endothelial transit determines the overall absorption for insulin lispr
- Insulin diffusion studied for the first time at the injection site of live animals
- *In vivo* imaging is a powerful tool to study injection site dynamics

## Introduction

Patients with insulin-dependent diabetes mellitus use rapid-acting insulins to control glucose excursions after meals. Most therapeutic insulins are available in stabilized hexamer forms, which after subcutaneous (SC) injection, dissociate into dimers and monomers for microvascular absorption, thus leading to a delay in time action [1–3]. Rapid-acting insulins such as insulin lispro (lispro) and insulin aspart are designed to have a higher hexamer-to-monomer dissociation rate that improves postprandial glucose control in patients; however, matching the rapid carbohydrate absorption profiles from meals still remains a challenge [3–6]. Both the aspart and lispro formulations use zinc and phenolic additives to form insulin hexamers [2,4]. While hexamer dissociation is widely considered to be rate-limiting for absorption of human insulin, research, to date, has not been able to confirm whether this is still the rate-limiting step for rapid-acting insulin analogues [7]. To produce even faster profiles, researchers have shifted their focus from modifications to the insulin molecule to changes in the drug product formulation to transiently alter the injection site tissue for enhanced microvascular absorption [8–10]. Understanding of insulin dynamics at the injection site is limited despite significant research in the field. Live visualization and quantification of insulin at the injection site can contribute to understanding critical processes, including injection site absorption kinetics and the effect of excipients on tissue properties. It has been nearly 100 years since insulin was first used therapeutically, yet there is no work in the literature characterizing insulin diffusion in a live animal. Previously, fluorescence, computed tomography (CT) scan, and isotope imaging have been used to study insulin boluses at the site of injection; however, those studies lacked live 3D structural and functional details covering the complete volume of injection at a cellular level resolution [11–13].

Photoacoustic microscopy (PAM) offers unique advantages over fluorescence and CT-scan imaging techniques. It can perform label-free imaging of blood and the measurement of local blood sO2 at a cellular level resolution [14,15]. PAM has never been demonstrated to quantify the absorption kinetics of biotherapeutics at the injection site because 3D quantitative imaging of an external contrast agent in live animals remains one of the significant challenges in the field of photoacoustic tomography [16].

In this article, we report the 3D qualitative and quantitative study of near-infrared (NIR) dye-labeled lispro in both the Humalog and Lyumjev formulations using optical-resolution photoacoustic microscopy (OR-PAM) in live mouse ear [17]. Up until now, there are no studies that detail mechanism of uptake of insulin bolus across the microvasculature at a cellular level resolution [18]. Using the murine ear cutaneous model lends itself to this kind of study due to its tractability, thinness, and compatibility with OR-PAM. While the aim of this study was not to equate murine ear skin to either pig or human skin subcutaneous layer, we anticipate that basic mechanisms of microvascular endothelial absorption of insulin should be conserved across those species particularly given the same organ (skin) in question and hence opening an avenue of new research investigation utilizing the murine ear to further interrogate mechanisms of biologic absorption and disposition. This lays the groundwork for studying the impact of extracellular matrix and vascular endothelium across tissues such as subcutaneous among different species. Insulin lispro in Humalog and Lyumjev drug products is formulated as a hexamer; however, the latter formulation contains treprostinil and sodium citrate to accelerate SC absorption [2,3,10]. To study absorption kinetics of the insulin formulations, we devised a novel 3D photoacoustic quantification approach for noise suppression. We modeled the experimental kinetics and diffusion data to understand the factors responsible for faster absorption in the Lyumjev formulation. We also carried out a study in pigs on the pharmacokinetics of unlabeled insulin lispro in Humalog, Lyumjev, and zinc-free, cresol-free formulations. We characterized the association state of insulin lispro in these formulations by sedimentation velocity analytical ultracentrifugation (SV-AUC) and 2D ^1^H-^13^C HSQC NMR. The results are consistent, across labeled and unlabeled insulin lispro, with a model in which crossing of the microvascular endothelium and not hexamer dissociation is the key rate-limiting step. This understanding paves the way for the development of improved insulin therapies for people living with diabetes.

## Materials and Methods

### Insulin dye labeling and characterization

#### Site-specific labeling of insulin lispro

The sulfo-cyanine7.5 (sulfo-cy7.5) labeled insulin analog was generated by selective acylation of the B-chain lysine B28 epsilon amino group of biosynthetic B28K, B29P-HI (insulin lispro), sourced from internal production, using a commercially available NHS ester from Lumiprobe. Insulin lispro zinc crystals (400 mg, 68.8 mmol) were dissolved (in a 20 mL scintillation vial) in 3 mL of 0.1 M sodium carbonate buffer (pH 10) along with 500 μL of acetonitrile to aid in dissolution. The vial was then placed on a magnetic stirrer with a suitable stir bar and the instrument was set to 300 RPM. To this solution are added 25 mg (21.2 mmol) of sulfo-cy7.5-NHS (46320, Lumiprobe) dissolved in acetonitrile (500 μL) in 100 μL portions over 10 min. The reaction was monitored by analytical LC-MS to follow the generation of the product. Once the reaction ceased (~ 1 h), the solution was diluted to 20 mL with ddH2O and loaded onto a preparative HPLC for immediate purification. RP-HPLC purification was performed on a Shimadzu LC-8A system equipped with a Rheodyne 3725i preparative sample injector (20 mL loop) and a semi-prep Luna 5 μm Phenyl-Hexyl RP-HPLC column (Phenomenex 00G-4257-P0-AX; 5 μm, 100Å; 2.1 x 25cm). The sample was eluted using a 15 – 40 % B gradient over 50 min at a flow rate of 25 mL/min at 60 °C (buffer A: 0.15% TFA in water and buffer B: acetonitrile). Fractions containing the desired product, (determined by analytical LCMS) were pooled, frozen, and lyophilized. The resulting TFA salt lyophilizate was then dissolved in 20 mL of 0.1 M NH_4_Cl (with ~ 5% acetonitrile) and subjected to another round of HPLC purification to perform a TFA to HCl salt conversion. Using the same system and column, the sample was loaded on a 10% B conditioned column, the gradient was held at 10% B for 10 min, the sample was then eluted using a 10 - 45 % B gradient over 70 min at a flow rate of 25 mL/min at 60 °C (Buffer A: 0.05% HCl in water and Buffer B: acetonitrile). Fractions containing the desired product, (determined by analytical LCMS) were pooled, frozen, and lyophilized to give 70 mg (HCl salt) of a green amorphous solid (48.9% of theoretical yield based on limiting reagent; 98% purity by RP-HPLC). The observed molecular weight was 6756.70 Da compared to a theoretical molecular weight of 6756.67 Da.

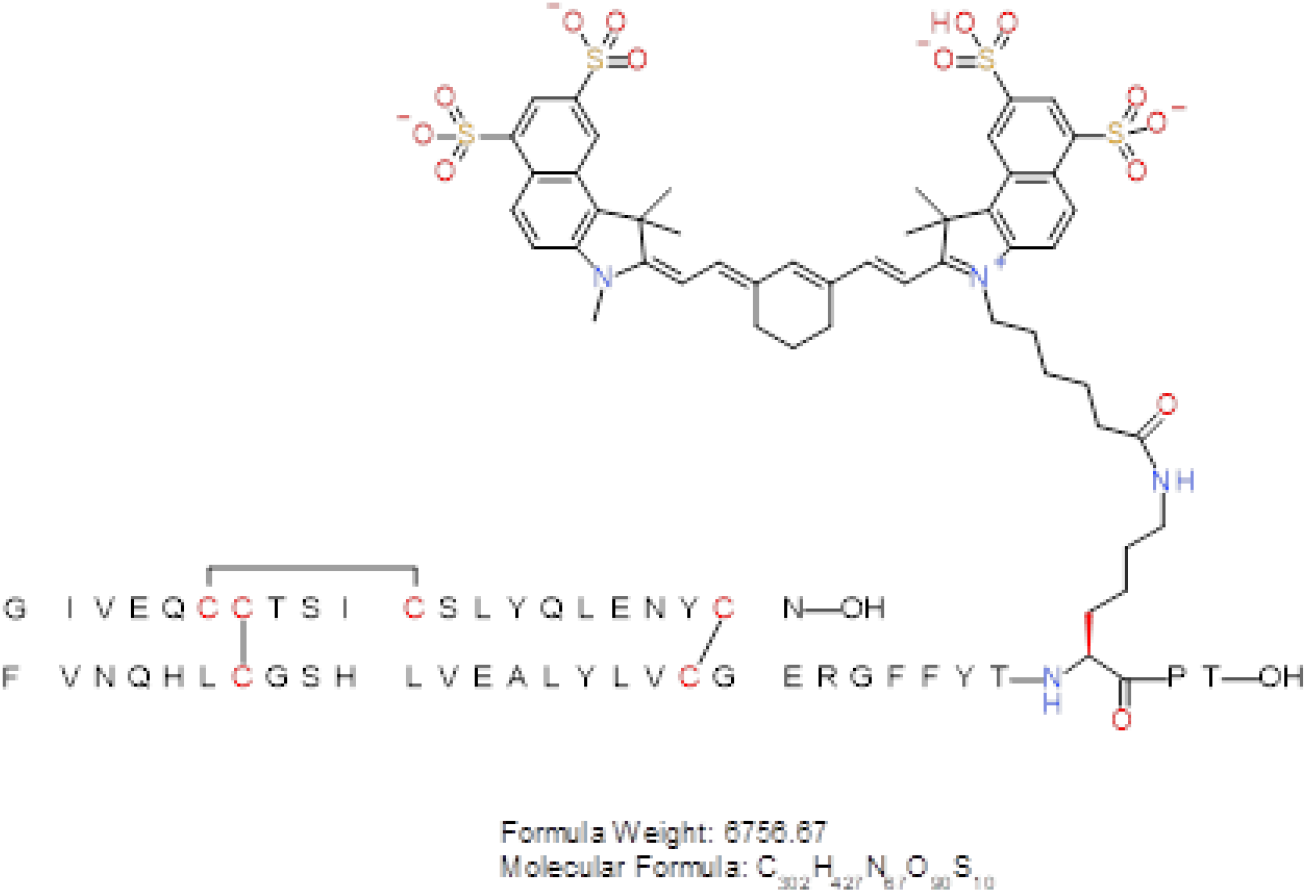

#### Lyumjev formulation

Lyophilized sulfo-cy7.5 insulin lispro was reconstituted in 20 mM NaOH to an approximate concentration of 16 mg/mL. Concentrated stocks of glycerol (400 mg/mL), m-cresol (20 mg/mL), zinc (30 mM in 100 mM HCl), sodium citrate (500 mM), magnesium chloride (500 mM), and treprostinil (1 mg/mL) were added in the order as mentioned. Deionized water was used to dilute to a final formulation composition of 1.2 mM sulfo-cy7.5 insulin lispro, 9 mg/mL glycerol, 3.15 mg/mL m-cresol, 0.8 mM zinc, 15 mM sodium citrate, 5 mM magnesium chloride, and 1 μg/mL treprostinil. The pH was adjusted to 7.4 using 1 N HCl. To prepare 400 μM sulfo-cy7.5 insulin lispro solution, the 1.2 mM sulfo-cy7.5 insulin lispro solution was diluted in the Lyumjev buffer solution (9 mg/mL glycerol, 3.15 mg/mL m-cresol, 0.8 mM zinc, 15 mM sodium citrate, 5 mM magnesium chloride, and 1 μg/mL treprostinil). The final solutions were filtered through a 0.2 μm pore membrane.

#### Humalog formulation

Lyophilized sulfo-cy7.5 insulin lispro was reconstituted in 20 mM NaOH to an approximate concentration of 10 mg/mL. Concentrated stocks of glycerol (400 mg/mL), m-cresol (20 mg/mL), zinc (30 mM in 100 mM HCl), and sodium phosphate dibasic (200 mM) were added in the order as mentioned. Deionized water was used to dilute to a final formulation composition of 1.2 mM sulfo-cy7.5 insulin lispro, 16 mg/mL glycerol, 3.15 mg/mL m-cresol, 0.6 mM zinc, and 7 mM sodium phosphate. The pH was adjusted to 7.4 using 1 N HCl. To prepare 400 μM sulfo-cy7.5 insulin lispro solution, the 1.2 mM sulfo-cy7.5 insulin lispro solution was diluted in the Humalog buffer solution (16 mg/mL glycerol, 3.15 mg/mL m-cresol, 0.6 mM zinc, and 7 mM sodium phosphate). The final solutions were filtered through a 0.2 μm pore membrane.

#### Sulfo-cy7.5 in Lyumjev, Humalog, and PBS buffer solutions

The sulfo-cy7.5 dye was dissolved in either Lyumjev or Humalog excipient formulations or PBS at concentrations of 1.2 mM or 400 μM. The concentrations of the solutions were matched by overlaying their Vis-NIR spectra.

### Measurement of the molar absorption coefficient of sulfo-cy7.5-insulin lispro in Humalog formulation

The sulfo-cy7.5 insulin lispro solution (1.2 mM) was diluted 300 times with the Humalog buffer solution and the absorbance was measured using a Varian Cary^®^ 50 Bio UV-Vis spectrophotometer at 20 °C. The molar absorption coefficient was calculated using Beer-Lambert law, A = ε.C.L where, A is the absorbance, ε is the molar absorption coefficient, C is the dye concentration, and L is the light path length (1 cm).

### Photoacoustic microscopy (PAM) system design

The photoacoustic microscopy (PAM) system employed a spherically focused ring-shaped transducer (central frequency = 42 MHz, f-number = 1.67, Capistrano Labs) placed confocally to the objective lens. The system uses light pulses of two optical wavelengths559 nm and 780 nm, which irradiate the same scanning pixel successively. The 559 nm light beam was focused for spatial filtering on a 254 μm diameter pinhole (3928T991, McMaster-Carr). The 780 nm light beam was generated using Styryl 11 dye (07980, Exciton) in 200 proof ethanol (MFCD00003568, Koptec) circulated in a dye laser (Credo, Sirah), which was pumped by a 532 nm Nd:YAG laser (IS80-2-L, Edgewave GmbH). The 780 nm light was then focused and spatially filtered through a second pinhole and then combined with the 559 nm light beam through a cold mirror (M254C45, Thorlabs). A separate pinhole for the 780 nm light beam was used to correct for the focal length difference and increase the depth of focus (~ 350 μm) for the 780 nm light beam. Both the combined beams were focused on the sample through an achromatic doublet (AC080-020-A, Thorlabs). A part of the light was directed to a photodiode (PDA36A, Thorlabs) by a beam sampler to correct for laser fluctuations. The raster scanning was performed by moving the animal using a stepper motor-based 2D scanner (PLS-85, Physik Instrumente GmbH). The scanning was controlled by an FPGA (PCI-e 7841, National Instruments) using a custom-designed program in LabVIEW. The photoacoustic and the photodiode data were acquired at 500 MS/s using a digitizer (ATS 9350, AlazarTech Inc.).

### Nano-liter injection

We used 34 gauge needles (207434, Hamilton) with microliter syringes to inject insulin volumes from 0.06 μL to 0.10 μL in the mouse ear. To perform the injection, the syringes were fitted onto a syringe dispenser, which enabled rapid and controlled injection. The needle was inserted in the mouse ear at an oblique angle (≤ 45°) and then the desired volume of the dye-labeled insulin lispro formulation was injected using the syringe dispenser. To perform 0.06 μL injections, a 1 μL syringe (80100, Hamilton) was used through which a minimum of 0.02 μL of liquid could be dispensed. To inject 0.1 μL volume, a 5 μL (7634-01, Hamilton) syringe was used. To perform 0.06 μL injection, the 1 μL syringe was fitted with a glass coupler (55752-01, Hamilton) and a luer adaptor (55753-01, Hamilton) to connect to the 34-gauge needle. The 5 μL syringe could be directly connected to the needle.

### Animal experiments

Mouse experiments were approved as per the IACUC regulations at Caltech. Hsd:Athymic Nude-Fox1^nu^ mice aged 6 to 10 weeks (Envigo) were used in all the photoacoustic experiments. The mouse body temperature was maintained at 37 °C during imaging. The mouse was continuously supplied with 1.25 – 1.50 % isoflurane in air to maintain anesthesia during imaging. The anesthetic state of the mouse was confirmed by pinching its hind paw.

Diabetic (alloxan induced), castrated, male Yucatan miniature swine with previously fitted vascular access ports were used for pharmacokinetic study under the supervision of staff and veterinarians at Sinclair Research Center (SRC), Auxvasse, MO. Treatment and Care of the animals are in accordance with regulations outlined in the USDA Welfare Act and conditions specified in “The Guide for Care and Use of Laboratory Animals.” All procedures involving the care or use of animals in this study were reviewed and approved by the SRC Institutional Animal Care and Use Committee prior to the initiation of such procedures.

Test articles were formulated at Eli Lilly and Company and shipped overnight on cold packs to SRC. They were stored refrigerated until the time of dosing and then returned to the refrigerator after the dosing of all animals was complete. During the dosing period, the test articles remained in an insulated box when not being withdrawn from.

The day prior to the study, animals were fed half their daily ration and received 0.2 U/kg Humalog Mix 75/25 Insulin at their morning maintenance administration. All study animals were food-fasted overnight and did not receive their evening insulin or meal prior to drug administration on the study day.

On the morning of the study, all animals were placed into slings for restraint and had their vascular access ports accessed (equipped for blood sampling) and checked for patency. The animals were randomly placed into treatment groups. After two baseline blood samples were collected (–30 and –20 min), the animals were returned to their pens and were fed ~300 g S-9 diet. Twenty minutes after the presentation of the fully consumed meal, the animals were injected with the test article subcutaneously (SC) in the flank (0 min) with a Terumo insulin syringe (0.3 or 0.5 ml with 1/2” needle). All study animals had *ad libitum* access to clean, fresh water throughout the remaining blood collection period.

This study was completed as a full crossover design where each animal was treated with each test article over three study dates.

Serial blood samples (2.0 mL each) were collected from each animal at the following time points: –30, –20 (then immediately fed), 0 (just before dose), 5, 10, 15, 30, 45, 60, 75, 90, 105, 120, 150, 180, 240, and 360 minutes following the SC dosing.

Blood samples without an anticoagulant were maintained at ambient temperature for at least 30 minutes but no more than 2 hours to allow for clotting. Serum was then separated by centrifugation and divided into two aliquots and stored frozen at approximately −70°C.

### Insulin formulations for pharmacokinetics

Zinc-free insulin lispro was produced by dialysis in 1000 MWCO membrane against four changes of phosphate buffered saline at 20X volume and three further changes of 0.1X phosphate buffered saline at 20X volume. Insulin lispro concentration after dialysis was 8.5 mg/ml and measured residual zinc content was 0.045 μg/mL. This material was formulated at 3.5 mg/mL with 16 mg/mL glycerol and 7 mM sodium phosphate and a final pH of 7.4. Hexameric lispro formulations approximating Humalog and Lyumjev were prepared at laboratory scale at U100 concentration. A 2x concentrated stock solution of insulin lispro zinc crystals, zinc (as ZnO), m-cresol, and glycerin was prepared in ultrapure water at pH ~ 7.4. For Humalog, sodium phosphate was added at the required amount to this 2x concentrated stock and then diluted with water to make up the final composition of 3.5 mg/mL insulin lispro, 3.15 mg/mL m-cresol, 12.1 mg/mL glycerin, 0.3 mM zinc, and 7 mM sodium phosphate. For Lyumjev, sodium citrate, treprostinil, additional zinc, and magnesium chloride hexahydrate were added to the 2x concentrated stock and then diluted with water to make up the final composition of 3.5 mg/mL insulin lispro, 3.15 mg/mL m-cresol, 12.1 mg/mL glycerin, 0.3 mM zinc, 1.0 mcg/mL treprostinil, 5 mM magnesium chloride hexahydrate, and 15 mM sodium citrate. Sodium hydroxide and hydrochloric acid were used to adjust pH as needed for all samples. All samples were analyzed by RP-HPLC and found to be within 5% of the intended insulin lispro concentration.

### Pharmacokinetic analysis

After extraction with polyethylene glycol (PEG), free serum insulin lispro concentrations were determined with an insulin lispro-specific sandwich ELISA using anti-insulin lispro antibody (Eli Lilly and Company) for capture and biotinylated anti-human insulin antibody (Eli Lilly and Company) with horseradish peroxidase conjugated streptavidin (7100-05, Southern Biotech) for detection. Insulin lispro standard curves and controls were prepared using Eli Lilly and Company insulin lispro reference standard in pig serum. The standard curve range was from 30 to 2400 pg/mL (5.2 to 413 pM) with a Lower Limit of Quantitation (LLOQ) value of 50 pg/mL (8.6 pM) and an Upper Limit of Quantitation (ULOQ) value of 2000 pg/mL (344 pM) (Charles River Laboratory, Quebec, Canada). Pharmacokinetic analysis was performed using Phoenix WinNonLin (version 6.4) and Microsoft Excel 2010, using a molar dose of 1.2 nmol/kg (0.2 U/kg). For analysis, samples below the LLOQ were assigned a value of 0 for timepoints prior to dosing, and 25 pg/mL (4.3 pM) for timepoints post-dose. Non-standard pharmacokinetic parameters to further explore shifts in the PK profile were also calculated and are defined in the relevant tables.

### Sedimentation velocity analytical ultracentrifugation (SV-AUC) analysis of Lyumjev, Humalog, and Monomeric Lispro

A Beckman Coulter ProteomeLab XL-I Analytical Ultracentrifuge was used for the study. The samples are placed in 3k cut-off centrifugal filters (Amicon Ultra −4 Centrifugal filters, Merck Millipore Ltd Cork, Ireland) and spun down at 4500 rpm for 25 minutes using a benchtop centrifuge. The flow through were used as a reference. 3 Beckman 2 sector cells equipped with sapphire windows were used for the study. The solvent and sample sectors of each cell were filled with 470 μL of reference and sample respectively and sealed with gaskets and screws. The cells were weighed, placed in a 4-hole rotor, and aligned using an optical alignment system. The rotor was loaded into the XL-I, allowed to temperature equilibrate to 20 °C for 2 hours, and the run was initiated at 60k rpm. Sedimentation velocity data was collected using Interference detection. SEDNTERP was used for individual density and viscosity values of the matrices [19]. The raw data were processed and fitted using SEDFIT program and continuous c(s) distribution [19]. The weight average sedimentation coefficients s(w) -values in the c(S) distribution were calculated for each sample run by integrating the distribution between 0.5 – 4 S values, the final reported weight average s(w) values are average of 3 independent sample runs. The insulin lispro concentration is the same across all samples at 3.5 mg/mL.

### NMR analysis

The NMR samples were prepared by mixing 540 μL of each sample with 60 μL of D_2_O. The resulting mixtures were transferred to 5-mm NMR tubes for analysis. All spectra were acquired at 25 °C on a Bruker Avance III 600 MHz NMR equipped with a 5-mm CP QCI H/F-C/N cryoprobe and with the hsqcetfpgpsi2 pulse sequence. The acquisition parameters included a 1-s relaxation delay, 0.1-s acquisition time, 14-ppm ^1^H sweep width, 30-ppm ^13^C sweep width, and 226 t1 increments (256 scans each). The resulting FIDs were zero-filled to 512 x 4096 points and weighted with cosine-squared functions in both dimensions prior to Fourier transformation. The baselines of the resulting spectra were corrected using the Whittaker Smoother algorithm in MestReNova (Mestrelab Research SL). The percent hexamer values were determined by comparing the volume of the hexamer crosspeak at *d_H_* 1.28*/d_c_* 18.99 with that of the non-hexamer crosspeak at *d_H_* 1.22/*d_c_* 19.50.

### Imaging protocol

For each insulin formulation, we imaged the mouse ear at a 4 kHz A-line scan rate with a fast axis (step size = 2.5 μm, 1100 steps) and a slow axis (step size = 5 μm, 800 steps), resulting in a total scanning time for a single image to be around 220 seconds. The following protocol was used for imaging the dye-labeled insulin lispro formulations in the mouse ear.

First, a pre-injection image of the mouse ear was taken. Following the first image, a sub-microliter injection of the required insulin formulation was performed in the mouse ear. The further images were taken at 3 minutes, 10 minutes, 15 minutes, 20 minutes, 25 minutes, 30 minutes, 45 minutes, 60 minutes, 90 minutes, 120 minutes, 150 minutes, and 180 minutes’ time points post-injection, during which the mouse was continuously under anesthesia. Following 180 minutes, the mouse was woken up and kept in its cage with food and water. The mouse was reimaged at 6 hours’ time point post-injection under anesthesia.

Since the time required for taking each image is around 220 seconds, we consider the mid-point of time at which the imaging was started and ended as the imaging time-point. For example, the scanning for the first image was started after 1 - 2 minutes post-injection, and it finished at 5 - 6 minutes post-injection. So, the time-point for the first image is assumed approximately at 3 minutes. For comparison of the kinetics data from different animals and different formulations, the first image taken at 3 minutes after injection was considered as the baseline, and the data from the rest of the images were divided by it to estimate the fraction of the formulations absorbed. All the statistical kinetics data reported in this study represent mean ± standard error of the mean unless otherwise mentioned. All the p values were calculated using unpaired two-sample t-tests. Shapiro-Wilk tests were done before performing the t-tests to confirm normality at a significance level of 0.05.

### 3D tissue expansion

In our 3D photoacoustic images, the lateral *x-y* information was derived from 2D scanning of the sample, and the depth *z* information was derived from the ultrasound (photoacoustic signal) detection time as *z* = *c·t* where, *c* is the speed of the ultrasound waves in biological tissue, and *t* is the time. Unipolar 3D images were generated by taking the signal envelope of the time domain (axial coordinate) using the absolute of the MATLAB-based Hilbert transform of the raw 3D photoacoustic data [20].

The temporal axial expansion of the dye-labeled insulin bolus was quantified to estimate tissue expansion in the axial direction (*z*-direction). The data was passed through a median filter in the *x-y-z* directions using a window of size 4 x 4 x 1 voxels. The size of each voxel is 2.5 μm (*x*-direction) x 5 μm (*y*-direction) x 3 μm (*z*-direction). The sharp top and bottom boundaries of the dye-labeled insulin bolus were estimated by thresholding the data by summation of the mean and six times the standard deviation of the background amplitude. We estimate the tissue expansion as the change of volume between the upper and lower boundaries of the bolus, which have common *x-y* coordinates for all the images. The choice of six times the standard deviation gives a continuous boundary for all the imaging time points (3 minutes to 45 minutes) and does not produce spurious voxels outside the dye-labeled insulin bolus. No temporal tissue movement took place in the lateral direction in the first 45 minutes, confirmed by co-registration of blood vessels.

### Calculation of the total amount of dye-labeled insulin through photoacoustic quantification Selection of the 3D region of interest

A rough 3D boundary around the volume occupied by the dye-labeled insulin bolus was manually selected, ensuring that all the photoacoustic signals from the dye-labeled insulin bolus are within the boundary. The 3D region of interest was then selected by applying a threshold level of the summation of the mean and three times the standard deviation of the background signal amplitude (after performing zero-mean across the time domain and taking absolute of the MATLAB-based Hilbert transform) on the volume occupied within the 3D boundary.

The total amount of the dye-labeled insulin was estimated as the sum of photoacoustic signal amplitudes in the 3D region of interest.

### Noise suppression

The detected signal consists of the photoacoustic signal from the dye-labeled insulin bolus and zero mean additive random thermal noise. Since the volume of the 3D region of interest comprises tens of thousands of voxels, the influence of the noise can be estimated in an average statistical sense. For each voxel *i*, the detected signal *v_i_* consists of photoacoustic signal *s_i_* and additive zero-mean thermal noise *n_i_* with a root mean square (RMS) noise voltage *ñ*. The RMS voltage 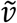 of *v_i_* can be expressed via signal and noise RMS powers:

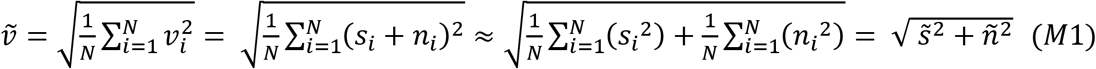

where *N* is the total number of voxels in the 3D region of interest and 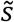 is RMS voltage of the ‘true’ signal. On average, the noise suppression can be performed by subtracting the noise power from the total detected power. However, because the concentration of the dye-labeled insulin is proportional to the true photoacoustic signal amplitude, we must add the true photoacoustic signal amplitudes rather than photoacoustic signal powers to calculate the total amount of the dye-labeled insulin.

The 3D region of interest was divided into several sub-regions such that the difference between the photoacoustic signal amplitude values of any two voxels in a sub-region is significantly smaller than the minimal photoacoustic signal amplitude within the same sub-region. For a sub-region *j*, the sum of the true photoacoustic signal amplitudes (*W_j_*) of all the voxels is estimated as:

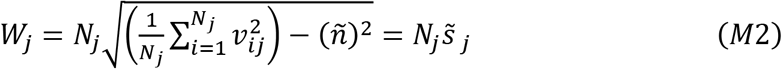

where *N_j_* is the total number of voxels in a sub-region.

Equation *M*2 gives a good approximation of the sum of the true photoacoustic signal amplitudes 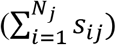 with an error on the order of 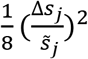, where Δ*S_j_* is the difference between the maximal and the minimal photoacoustic signal amplitudes in a sub-region. Following the calculation of summation of the true photoacoustic signal amplitudes of all the sub-regions, they are added together to estimate the total amount of the dye-labeled insulin in the bolus.

### Kinetic model

The kinetic model is based on compartmental modeling [21,22]. The model assumes that the hexameric insulin with initial concentration, *C_inj_* injected in the mouse ear tissue dissociates into monomers and transports in the interstitium at a rate constant *k*_1_ before reaching an equilibrium concentration, *C_eq_* (Methods). The insulin is then absorbed by the vasculature system at a rate constant *k*_2_. The equations *M*3 and *M*4 denote the relationships between, *C_inj_*, *C_eq_*, *k*_1_, and *k*_2_.

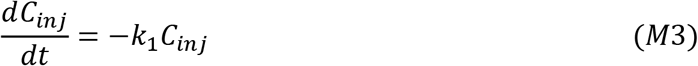

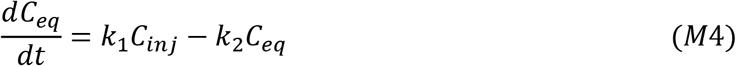

Since we are imaging *C_inj_* + *C_eq_*, we solved the above two differential equations to form equation *M*5,

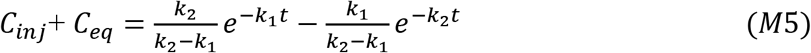

Moreover, a constant term *C*1 was added to equation *M*5 to account for the insulin precipitation/retention at the injection site. The boundary conditions for solving the above equations assume that at *t* = 0, *C_inj_* = 1 and *C_eq_* = 0. The term *C*1 was added to equation *M*5 such that at *t* = ∞, *C_inj_* + *C_eq_* = *C*1. A proportionality constant, *C* was finally multiplied to form equation *M*6 before fitting on the mean of experimental data; since the photoacoustic amplitude is linearly proportional to the dye-labeled insulin concentration.

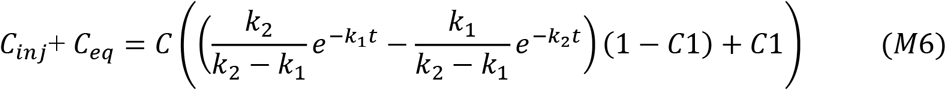

The fittings were performed by minimizing the sum of squared errors (SSE) in the nonlinear least-square sense. The values of *k*_1_ and *k*_2_ were estimated by fixing the lower bounds for *C, C*1, *k*_1_, *k*_2_ at 0. The starting points were set as *C* = 1, *C*1 = 0.03, *k*_1_ = 0.01, *k*_2_ = 0.011.

### Diffusion coefficient calculation

Trajanoski *et al.* calculated diffusion coefficients through simulation by solving equation 1 (in the main text) assuming that the insulin bolus distributes in a spherical geometry after injection [23]. In our case, we assume that the dye-labeled insulin lispro is spread in a cylindrical geometry because of the narrow thickness of the mouse ear and how the insulin was found to be distributed across the axial direction.

Upon considering a cylindrical geometry, equation 1 can be rewritten as equation *M*7

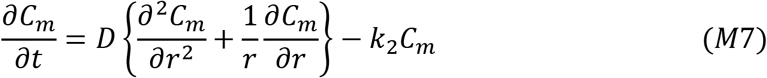

Equation *M7* has a closed form solution represented by equation *M*8 [24]:

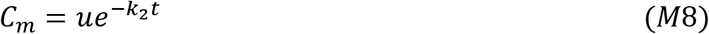

where:

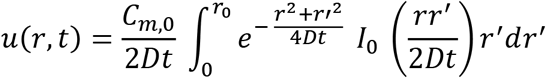

Here, *I*_0_ is modified Bessel function of the first kind, *r* is the distance from the center to the point of interest, *C*_*m*,0_ is initial dye concentration, *r*_0_ is the radius of the area occupied by the insulin at time *t* = 0, *k*_2_ is the vasculature absorption rate of the dye-labeled insulin (disappearance rate from the injection site), and *D* is the diffusion coefficient.

The lateral area covered by the dye-labeled insulin bolus changes temporally as per the diffusion and can be expressed in terms of *r*. Hence, by estimating the temporal change of the area, the diffusion coefficient *D* can be calculated.

Injection doses (n = 3) with higher dye-labeled insulin concentration (0.1 μL, 1.2 mM) were used for the first 60 minutes for an improved contrast-to-noise ratio at the boundaries. The raw photoacoustic data was converted to maximum amplitude projection (MAP) images to calculate the lateral area (in terms of *r*) occupied by the dye-labeled insulin bolus. The area can be estimated by thresholding the concentration of the dye-labeled insulin up to a fraction of its maximum value (assumed to be at the center of the bolus). Before estimating the area by taking a fraction of the maximum concentration, the MAP images were thresholded (after passing through a median filter of size 4 x 4 pixels) by the summation of the mean and three times the standard deviation of the background amplitude to segregate photoacoustic signals (from the dye-labeled insulin bolus) from the noise. Then the final area was selected by further thresholding the images with the fractional value of 0.1 of the average of the first 10 pixels with the maximum intensity. The area of the injection site was excluded from the search for the pixels with the maximum intensities. Due to the nature of equation *M8*, the smaller the fraction of the maximum concentration used as a thresholding value, the more sensitive is insulin area to the value of *D*, and the more accurate is the estimation of *D*. However, in practical terms, the area calculated by the fraction of the maximum concentration should be smaller or equal to the area estimated by the threshold of the summation of mean and three times the standard deviation of the background. We found that the value of 0.1 as the fraction of maximum concentration is an optimal choice to satisfy the above criteria in all our experiments.

The value of *D* was calculated by minimizing the difference between experimental temporal area change data and a similar theoretical solution of equation *M*8 in a nonlinear least square sense. Since we do not have area data at 0 minutes as required for the model, it serves as another unknown parameter. For all the cases, the area at zero time was found to be in the range of 0.83 to 0.95 of the area at 3 minutes. Note that as per equation *M*8, the radial distribution of concentration of the dye-labeled insulin as a function of time does not depend on *k*_2_. While fitting the experimental data, we exclude the first time point and used the time-points from 10 minutes to 60 minutes because in the image taken just after injection (i.e. at 3 minutes), the shape of the insulin bolus was driven by the injection pressure rather than diffusion.

## Results

### Dye-labeling of the lispro

We labeled the lispro molecule site-selectively with a photostable NIR light-absorbing sulfo-cyanine7.5 (sulfo-cy7.5) dye at the LysB28 position. We formulated the lyophilized dye-labeled lispro in either Humalog or Lyumjev formulation at concentrations of 1.2 mM and 400 μM corresponding to drug product concentrations of 200 units/mL and 67 units/mL.

We chose to study the dynamics of dye-labeled lispro formulations in a mouse ear model because of the thinness of the tissue (~ 200 - 250 μm), which minimizes tissue-based light-scattering and absorption of the 780 nm near-infrared light, thereby allowing reliable 3D quantification of external contrast agents.

### OR-PAM design and imaging parameters

To monitor the dynamics of the dye-labeled lispro (Figure 1a), we employed a custom-designed dual-wavelength (559 nm and 780 nm) OR-PAM (Figure 1b). We used the 559 nm wavelength to image blood label-free and the 780 nm wavelength to image the sulfo-cy7.5 dye-labeled lispro (Figure 1c).

**Figure 1:**
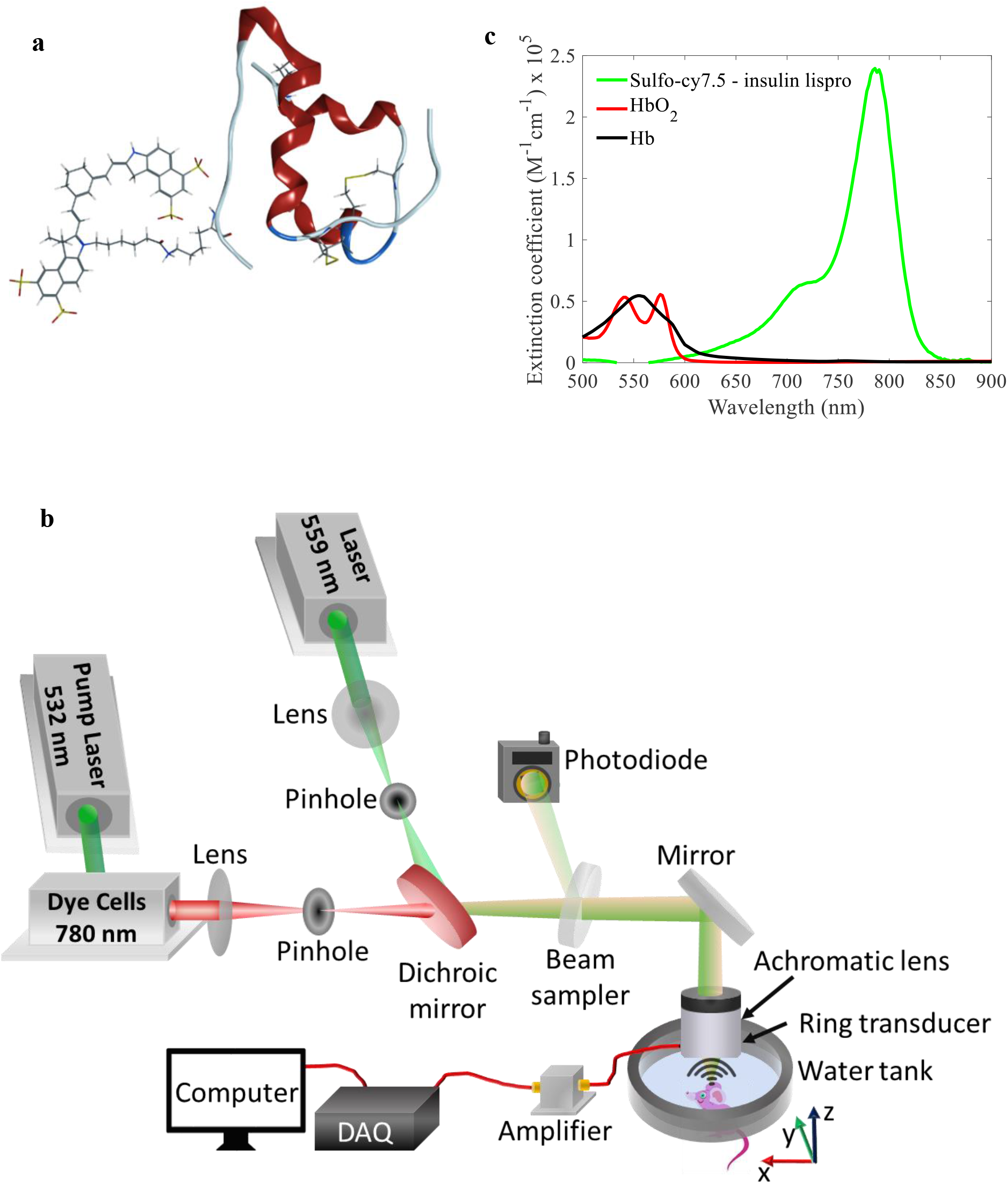
(a) Insulin lispro (lispro) with sulfo-cy7.5 dye covalently attached to the epsilon amine of LysB28. (b) Custom-designed OR-PAM. (c) Vis-NIR light absorption spectra of sulfo-cy7.5 dye-labeled lispro, oxygenated blood (HbO_2_), and deoxygenated blood (Hb). The values of blood absorption spectra were adopted from a compilation of Scott Prahl using data from N. Kollias, Wellman Laboratories, Harvard Medical School, Boston and W. B. Gratzer, Med. Res. Council Labs, Holly Hill, London.

We injected the insulin formulations at volumes ranging from 0.06 μL to 0.10 μL. Low injection volume enabled the use of clinically relevant concentrations of lispro (U67 and U200) without incurring hypoglycemia [25,26].

Before performing live animal experiments, we injected the dye-labeled lispro formulation in the ear of an expired mouse to study the progression of photoacoustic signals to assess photobleaching. We confirmed the photostability of the dye-labeled lispro under our imaging parameters by observing no decrease in photoacoustic signals after 21 imaging sessions (Figure S1).

### Live animal imaging

We injected dye-labeled lispro prepared in either the Humalog or Lyumjev formulation (1.2 mM, 0.10 μL, 5 nmol/kg in a ~25 g mouse) in the ear of an anesthetized mouse and then performed photoacoustic imaging. The local lymphatic vessels absorbed a portion of the dye-labeled lispro in both formulations immediately after injection due to the interstitial stress generated by the microneedle injection pressure in the mouse ear [27,28]. We saw that the blood microvascular system gradually absorbed the majority of the dye-labeled lispro in both the formulations (Figures 2a and 2b, movies S1 and S2).

**Figure 2:**
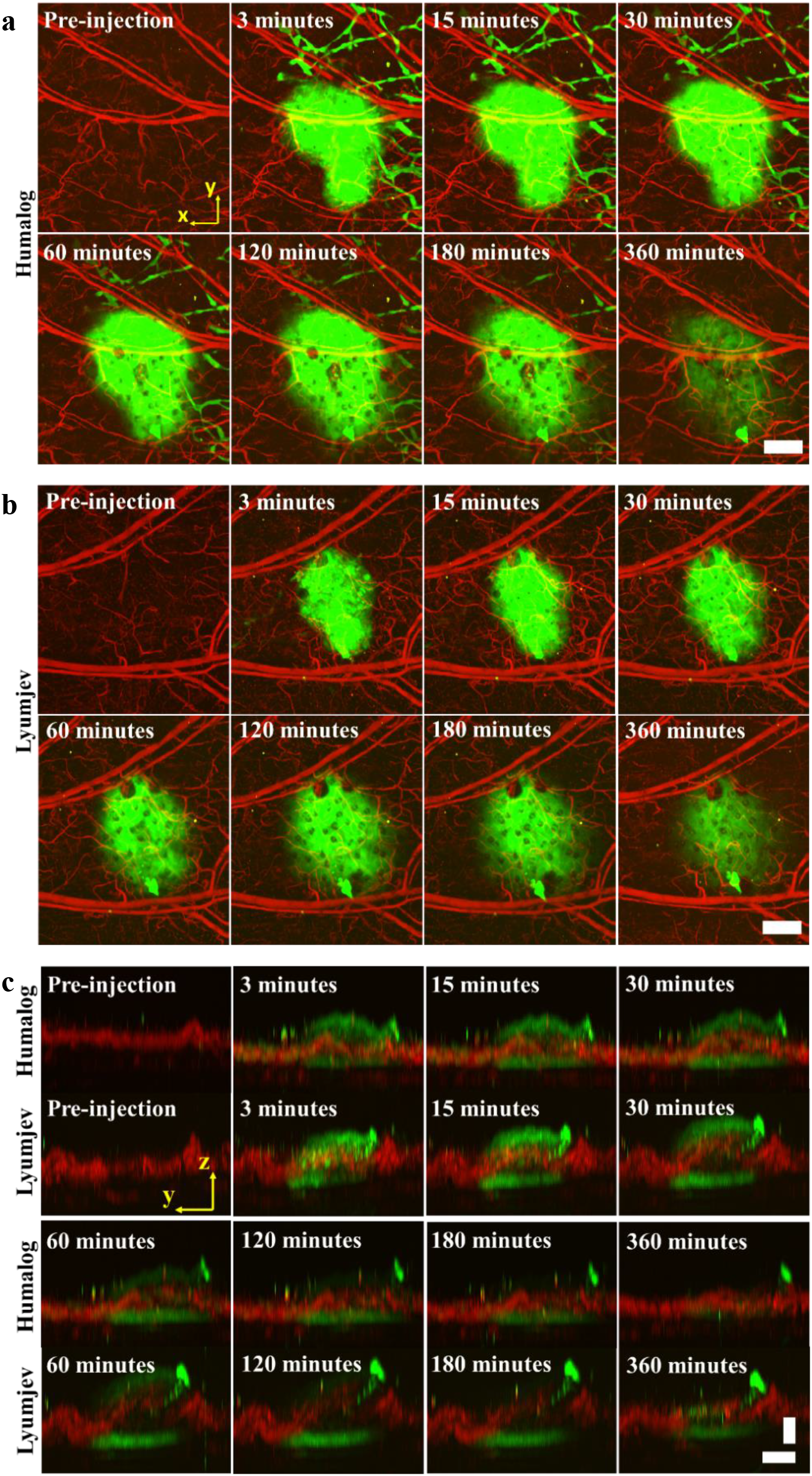
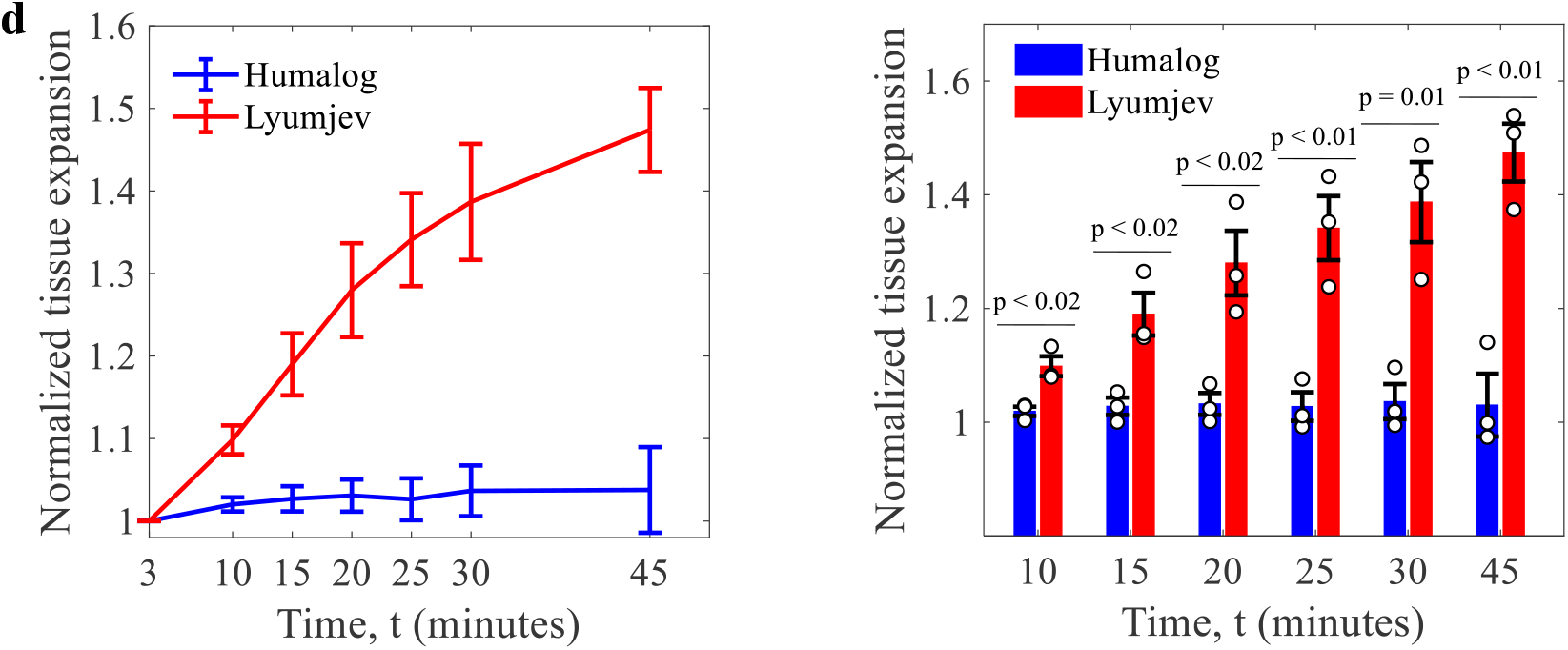
Visualization of dynamics of the insulin formulations in the mouse ear. Absorption of sulfo-cy7.5 dye-labeled lispro in either Humalog (a) and Lyumjev (b) formulations in mouse ear. (c) Side-view images of the sulfo-cy7.5 dye-labeled lispro in Humalog and Lyumjev formulations depicting tissue expansion. (d) Comparison of expansions caused by the Humalog and Lyumjev formulations, n = 3. Vertical scale bar = 100 μm; horizontal scale bar = 500 μm. Green is the sulfo-cy7.5 dye-labeled lispro and red is the blood (label-free). Data represent mean ± standard error of the mean.

As expected, the insulin formulations produced dye contrast in the lateral directions that exhibited a smooth decay, corresponding to injection site diffusion (Figures 2a and 2b); however, the boundaries in the axial direction presented a sharp demarcation of a moving top boundary, indicating tissue expansion (Figure 2c). The temporal change in the morphology of blood vessels observed in the axial direction, due to the Lyumjev formulation, indicates modification of the injection site by the added excipients. We measured the mean expansion induced by the Lyumjev formulation after 45 minutes at ~ 1.5x compared to ~ 1.0x induced by the Humalog formulation (Figure 2d). No tissue movement took place in the lateral direction in the first 45 minutes, as confirmed by co-registration of blood vessels.

### Photoacoustic quantification of lispro

We quantified the absorption kinetics using OR-PAM to better understand how the differences in the two formulations affect the absorption of the dye-labeled lispro. Photoacoustic signal amplitude generated from the dye-labeled lispro is directly proportional to the concentration; hence, the sum of photoacoustic signal amplitudes from the entire dye-labeled lispro bolus is an estimate of the total amount (Figure S2) [16]. To estimate the total amount of the dye-labeled insulin, we summed the photoacoustic speckle amplitudes in the bolus after performing noise suppression (see Methods). We performed imaging at a reduced injection dosage (400 μM, 0.06 μL) to ensure an optically thin sample to minimize absorber-based light attenuation [29].

We confirmed the accuracy of quantification by observing linear proportionality between the injected doses (at equal volumes of 0.06 μL), and measurement of total amounts of dye-labeled lispro in an expired mouse-ear (Figure 3a).

**Figure 3:**
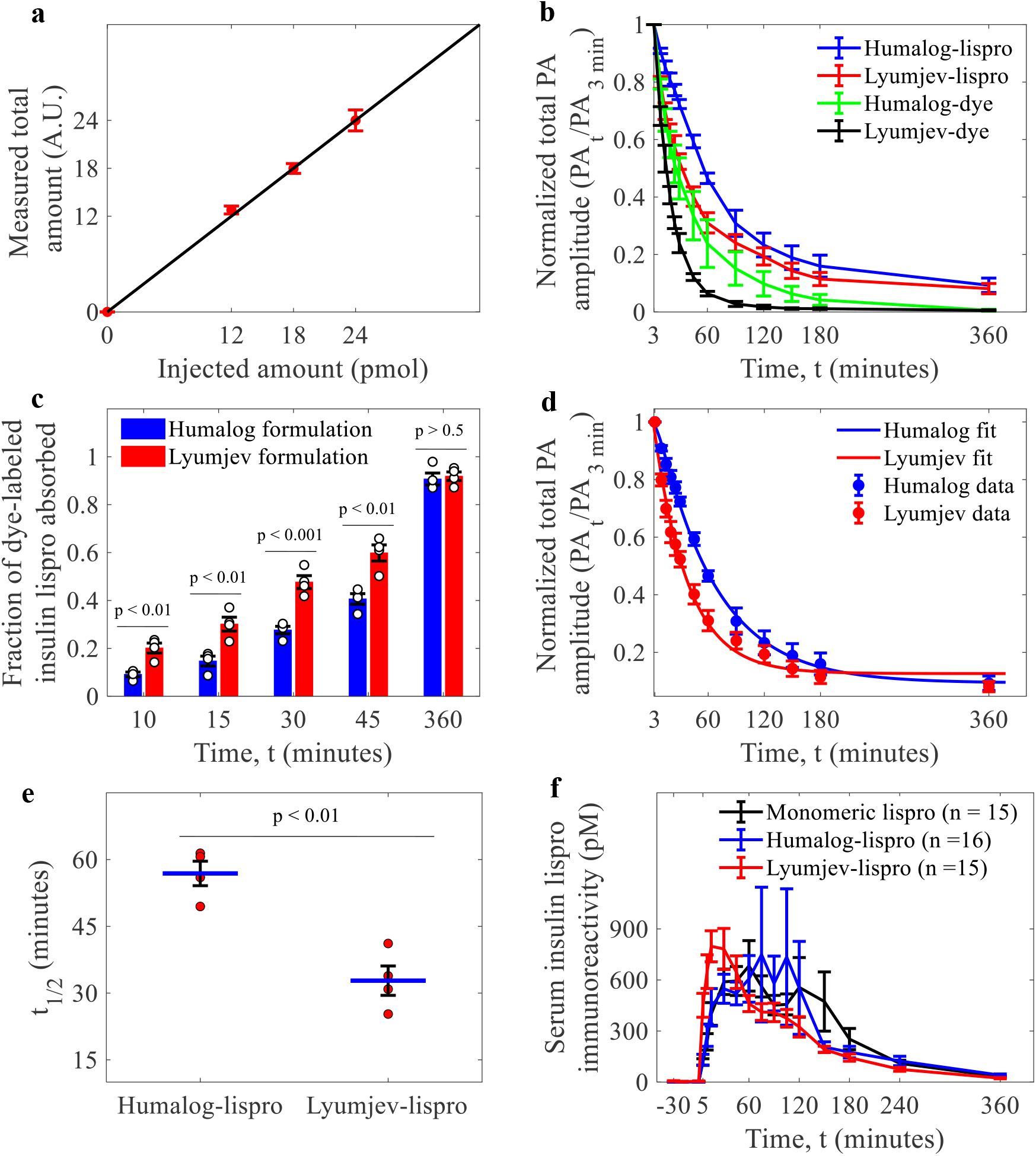
Photoacoustic (PA) quantification and pharmacokinetics study of the insulin formulations. (a) Linear dependence of the measured total amounts of the dye-labeled lispro on their injected amounts. Data (n = 3) represent mean ± standard deviation. (b) Comparison of injection-site absorption kinetics of the dye-labeled lispro (n = 4) and sulfo-cy7.5 dye (n = 3) in the Humalog and Lyumjev formulations. (c) Comparison of the fractional injection-site absorption for dye-labeled lispro in both formulations; n = 4. (d) Kinetic model fit on the mean of the absorption for dye-labeled lispro in both formulations; n = 4. (e) Comparison of t_1/2_ (time required for 50% drug absorption). (f) Pharmacokinetic study in pigs for monomeric lispro (n = 15) in Humalog formulation without zinc and phenolic ligand, hexameric lispro (n = 16) in Humalog formulation, and hexameric lispro (n = 15) in Lyumjev formulation. Data represent mean ± standard error of the mean.

We injected the dye-labeled lispro in both the Humalog and Lyumjev formulations (0.06, 1 nmol/kg in a ~25 g mouse, U67, Figure S3) and sulfo-cy7.5 dye without lispro, in the same formulations, in the mouse ear and studied the absorption profiles over 6 hours (Figure 3b).

The Lyumjev formulation resulted in faster absorption of the dye-labeled lispro than the Humalog formulation. We compared the fractional absorptions at different time points to assess the temporal effect of the excipients in the Lyumjev formulation (Figure 3c). Within the initial 15 minutes of imaging, the amount of dye-labeled lispro absorbed was 2.0 - 2.5x greater in the Lyumjev formulation compared to the Humalog formulation. Subsequently, over the first 45 minutes, dye-labeled lispro absorbed from the Lyumjev formulation was consistently greater than that observed in the Humalog formulation, while no differences were observed after 6 hours (Figure 3c). Since citrate and treprostinil resulted in faster absorption of lispro in the Lyumjev formulation, we compared the absorption of just the sulfo-cy7.5 dye in each formulation (Figure 3b). As expected, the sulfo-cy7.5 dye was absorbed at a faster rate in the Lyumjev formulation (Figures 3b and S4a). Upon comparing the absorption of the dye-labeled lispro with that of sulfo-cy7.5 dye in the respective formulations, we observed that the overall absorption of the dye-labeled lispro was slower than the dye (Figures S4b and S4c).

### Kinetic modeling and pharmacokinetics

The absorption of the dye-labeled lispro from the injection site is dependent on different factors; thus, for better understanding, we adapted a compartmental model designed for rapid-acting insulins for our imaging experiments to determine the rate constants, *k*_1_ and *k*_2_ (see Methods) [21,22]. The rate constant *k*_1_ includes both the rate of hexamer-to-monomer dissociation and lispro diffusion in the interstitium while *k*_2_ refers to the rate of lispro absorption across the vasculature.

We report the values of *K*_1_(Table 1), at the minimum SSE, to estimate the order of magnitude for the dye-labeled lispro dissociation and diffusion in the Lyumjev and Humalog formulations (Figure 3d). Notably, *k*_1_ cannot be precisely determined due to the rapidity of the two physical processes; thus, we report the values at the minimum SSE. Nonetheless, the model can tightly estimate *k*_2_, and since *k*_1_ » *k*_2_ (Table 1), we can conclude that absorption of insulin lispro across the microvascular endothelium is the rate-limiting step. Upon comparing the mean microvascular absorption rate constants, of the two formulations, we found that the rate constant associated with the Lyumjev formulation is ~1.7x greater than in the Humalog formulation.

**Table 1:** Rate constants of dye-labeled lispro in the formulations

| Compound | $k_1$ (min<sup>-1</sup>) | $k_2$ (min<sup>-1</sup>), 95% confidence bounds |
| --- | --- | --- |
| Humalog - lispro | 0.2 | 0.016 (0.014 – 0.018) |
| Lyumjev - lispro | 2.8 | 0.027 (0.021 – 0.034) |

From modeling the experimental data, we calculated *t*_1/2_ (i.e., time to 50% absorption) of the dye-labeled lispro in the Lyumjev formulation at 32.8 minutes, which was almost half of that in the Humalog formulation at 56.9 minutes (Figure 3e).

We studied the pharmacokinetics of unlabeled lispro in Humalog, Lyumjev, and predominantly monomeric zinc-free, cresol-free formulations in pigs (Fig 3f, Fig S5, and Table S1) to confirm our assessment that hexamer dissociation rate is not the absorption rate-limiting step. The overlap of both the pharmacokinetic profiles, especially in the first 0 – 30 minutes shows that hexamer dissociation of lispro in Humalog is too fast to be the rate-limiting step for absorption. The lispro in the Lyumjev formulation is absorbed ~1.8x faster than in the Humalog formulation in the first 30 minutes, which is consistent with the photoacoustic measurements of the dye-labeled lispro.

### Association state for unlabeled formulations

To confirm that Humalog and Lyumjev formulations are hexameric and to characterize the association state of the zinc-free, cresol-free formulation, we carried out sedimentation velocity analytical ultracentrifugation and NMR methyl ^1^H-^13^C HSQC studies. Sedimentation of insulin lispro in Humalog and Lyumjev formulations was consistent with a hexamer, while sedimentation in the zinc- and cresol-free formulation was consistent with a predominant monomer (Table 2).

**Table 2:** Weight average sedimentation coefficient s(w) values.

| Formulation | $s(w)$ | $SD$ |
| --- | --- | --- |
| Humalog | 2.83 | 0.002 |
| Lyumjev | 2.90 | 0.001 |
| Zinc-free, cresol-free | 1.22 | 0.007 |

Hexameric and monomeric forms of insulin lispro exhibited significantly different ^1^H-^13^C HSQC spectra (Figure S6), as expected based on different chemical environments for residues in the hexamer interfaces. We integrated sampling of resonances unique to each form to characterize the equilibrium between monomer and hexamer for each formulation and found that the Humalog and Lyumjev formulations were almost completely hexameric, with only 7% and 2% respectively in the monomeric form (Table 3). On the other hand, no hexamer was detected in the zinc-free, cresol-free formulation.

**Table 3:** Association state equilibrium by ^1^H-^13^C HSQC.

| **Formulation** | ***Hexameric*** | ***Non-hexameric*** |
| --- | --- | --- |
| Humalog | 93% | 7% |
| Lyumjev | 98% | 2% |
| Zinc-free, cresol-free | 0% | 100% |

### Diffusion

We determined the diffusion coefficients of dye-labeled lispro in the Humalog and Lyumjev formulations by analyzing the temporal change in the 2D area covered by the compounds in the mouse ear to characterize the diffusion. We calculated the diffusion coefficients by fitting the temporal area change data through a mathematical model that we derived from diffusion equation 1 (see Methods) [23].

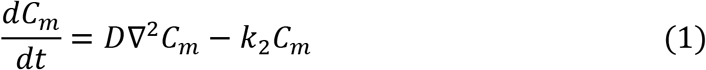

wherein, *C_m_* is the concentration of monomers, *D* is the diffusion coefficient, ∇ is the Laplacian operator, *k*_2_ is the absorption rate constant, and *t* is time. As evident from the pharmacokinetics study, the hexamer dissociation is complete in the first few minutes after injection. We report the diffusion coefficients (Table 4) for dye-labeled lispro monomers derived from equation 1 by fitting the model on the temporal change of the area of the insulin bolus from 10 minutes to 60 minutes post-injection (Figure S7). We found the mean diffusion coefficient of the dye-labeled lispro to be higher in the Lyumjev formulation.

**Table 4:** Diffusion coefficients of the dye-labeled lispro in the formulations.

| Compound | $D \text{ (cm}^2\text{/s)} \pm \text{S.D.}$ | $D \text{ (}\mu\text{m}^2\text{/min)} \pm \text{S.D.}$ |
| --- | --- | --- |
| Humalog - lispro | $1.84 \times 10^{-8} \pm 0.4 \times 10^{-8}$ | $110 \pm 24$ |
| Lyumjev - lispro | $7.54 \times 10^{-8} \pm 2.2 \times 10^{-8}$ | $452 \pm 132$ |

Our imaging shows that the blood vessels are spaced at a distance of about 5 – 20 μm. Hence, assuming a similar diffusion distance of insulin post-injection, we calculated the diffusion rate constants for dye-labeled lispro in Lyumjev and Humalog formulations and found them to be on the order of 2 min^-1^ and 0.5 min^-1^ respectively. Since the values of *k*_1_ for the compartmental model estimate similar orders of magnitude for the combination of hexamer dissociation and diffusion, we conclude that *k*_1_ is dominated by diffusion, and hexamer dissociation for dye-labeled lispro is too fast to make a significant contribution to the rate constant.

## Discussion

We used photoacoustic imaging to study injection-site absorption kinetics of dye-labeled lispro in both the Humalog and Lyumjev formulations in murine ear skin and found that dye-labeled lispro is absorbed more than twice as fast in the latter formulation in the first 10 to 15 minutes. This is in agreement with our pharmacokinetic analysis in pigs as well as the reported clinical trials data [10,30]. Our results show that the Lyumjev formulation is associated with local tissue expansion, faster diffusion of labeled insulin in the interstitium, and faster absorption into the blood vessels. Increases in the permeability of the microvascular endothelium and the extracellular matrix, along with the observed tissue expansion, are likely to be the consequence of chelation of calcium by citrate while treprostinil produces localized vasodilation at the injection site [31–33].

Our pharmacokinetics data on pigs along with the kinetics and diffusion modeling of the experimental data from mice show that hexamer dissociation rate and diffusion are not the rate-limiting step for dye-labeled lispro absorption from either the Humalog or Lyumjev formulation. While we found that *k*_2_, representing endothelial transit, was rate-limiting for the overall absorption, *k*_1_, representing diffusion through the extracellular matrix, can still make meaningful contributions to the early absorption, i.e., around the first minute. This is important since the appearance of insulin lispro in systemic circulation in clinical studies is reported to happen at 1.26 minutes and 6.84 minutes for the Lyumjev and Humalog formulations, respectively [10]. Modifying the permeability of the extracellular matrix and/or vascular endothelium has previously been demonstrated to be sufficient to speed insulin absorption in studies using hyaluronidase [34]. Literature accounts on the hexamer dissociation rates for rapid-acting insulins are scarce and contradictory [7,22]. Though it has been suggested, without convincing proof, that hexamer dissociation rate for lispro is the rate-limiting step in absorption [22], these results show that absorption across the microvasculature plays a more significant role. A previous study to characterize the difference in pharmacokinetics between hexameric and non-hexameric formulations of lispro in pigs was compromised by the presence of phenolic preservative in the absence of zinc, which results in higher order association [3]. To get a clearer understanding of the effect of hexamerization on pharmacokinetics, we compared a fully stabilized hexameric formulation containing zinc and cresol with the preparation of lispro lacking both, which we have demonstrated exhibits a predominantly monomeric association state [3,35,36]. Pharmacokinetics of these two formulations are indistinguishable from 5 to 60 minutes, indicating that hexamer dissociation is rapid relative to that timeframe and not rate-limiting for absorption. Studies on cobalt (III) stabilized hexamers indicate that hexamers cross the microvascular endothelium slowly and direct absorption of hexamers is too slow to contribute significantly to absorption on this timescale [37,38]. The findings reported here indicate that in the field of rapid-acting insulin research, focusing on hexamer dissociation rates of rapid-acting insulins may not be the optimal strategy. Since the overall absorption of dye-labeled lispro in the Lyumjev formulation is limited by the rate at which it crosses the microvascular endothelium (*K*_2_), the next logical approach to create even faster insulins is to focus on transiently increasing the rate of vascular absorption. Moreover, the *in vivo* diffusion analysis shows that excipients do influence protein transport by enabling faster diffusion in the interstitium. Until now, all the reported studies on diffusion of therapeutic insulins have been performed in either water, gel-based phantoms, or *ex vivo* [25,39,40]. Our *in vivo* study accounts for crucial diffusion factors, such as interstitium pressure on the drug from tissue elasticity, the effect of interstitial fluid viscosity, and tissue heterogeneity that were not properly controlled for in the past studies and suggests a further study to understand the translation of this mouse ear cutaneous model to the human subcutaneous tissue. Our diffusion model based on fitting the area of the bolus, can be applied to a wide range of shallow (< 1 mm) and deep (> 1 mm) imaging techniques for different types of injectable drugs.

In summary, we demonstrated the use of photoacoustic imaging to qualitatively and quantitatively study the injection site dynamics of rapid- and ultra-rapid insulin formulations. We addressed critical questions on the mechanisms of enhanced absorption of the Lyumjev formulation, including microvascular transport, drug diffusion, and tissue expansion. These experiments coupled with the kinetic model show that hexamerization of rapid-acting insulin analogs and subsequent hexameric dissociation rate for absorption is not the rate-limiting step for lispro, which has been a question of debate for more than 20 years. Overall, the application of OR-PAM demonstrates that the methodology can play a significant role in understanding the real-time behavior of a wide range of therapeutics at the injection site.

## Supporting information

Supplementary Images

## Acknowledgments

We thank Steven Cottle and Linh Nguyen, Eli Lilly and Company for preparing the insulin formulation solutions. We also thank Jake Anderson, Eli Lilly and Company for useful discussions. This work was funded by Eli Lilly and Company.

## Data availability

The data that support the conclusions are present in the main draft or the supplementary information. The data processing and quantification procedures are described in detail in the Methods section.

## Author contributions

L.V.W., S.O., J.M.B., C.D.P., and A.K. conceived the project and the ideas. C.D.P. designed the chemistry for dye labeling and analyzed the chemistry data. F.A.V. labeled the insulin molecules with the sulfo-cyanine7.5 dye and characterized them. F.A.V. and A.K. prepared the insulin and dye buffer solutions. A.K. and K.M. designed and built the scanning photoacoustic microscope. J.S. and R.C. wrote the LabVIEW software for photoacoustic data acquisition. A.K. designed and performed all the photoacoustic experiments, wrote the MATLAB codes, and analyzed all the photoacoustic data. A.K., K.M., and L.V.W. designed the photoacoustic quantification algorithm. A.K. and K.M. devised the pharmacokinetics and diffusion methodologies. A.L.C. designed and supported the pharmacokinetics study in pigs and A.E.S. performed the pharmacokinetic experiments. R.M. prepared the unlabeled insulin lispro samples, and designed and performed the SV-AUC experiments. S.A.B. designed and performed the NMR spectroscopy analyses. A.K., C.D.P., K.M., J.M.B, and L.V.W. interpreted the final data. P.B.A., E.L., and R.L.B. were involved in active discussions. L.V.W., S.O., and J.M.B. supervised the project. A.K. wrote the manuscript. C.D.P., K.M., R.C., J.M.B, S.O., and L.V.W. contributed to writing the manuscript.

