## Supplementary Images for "Photoacoustic imaging reveals mechanisms of rapid-acting insulin formulations dynamics at the injection site"

### Supplementary Text

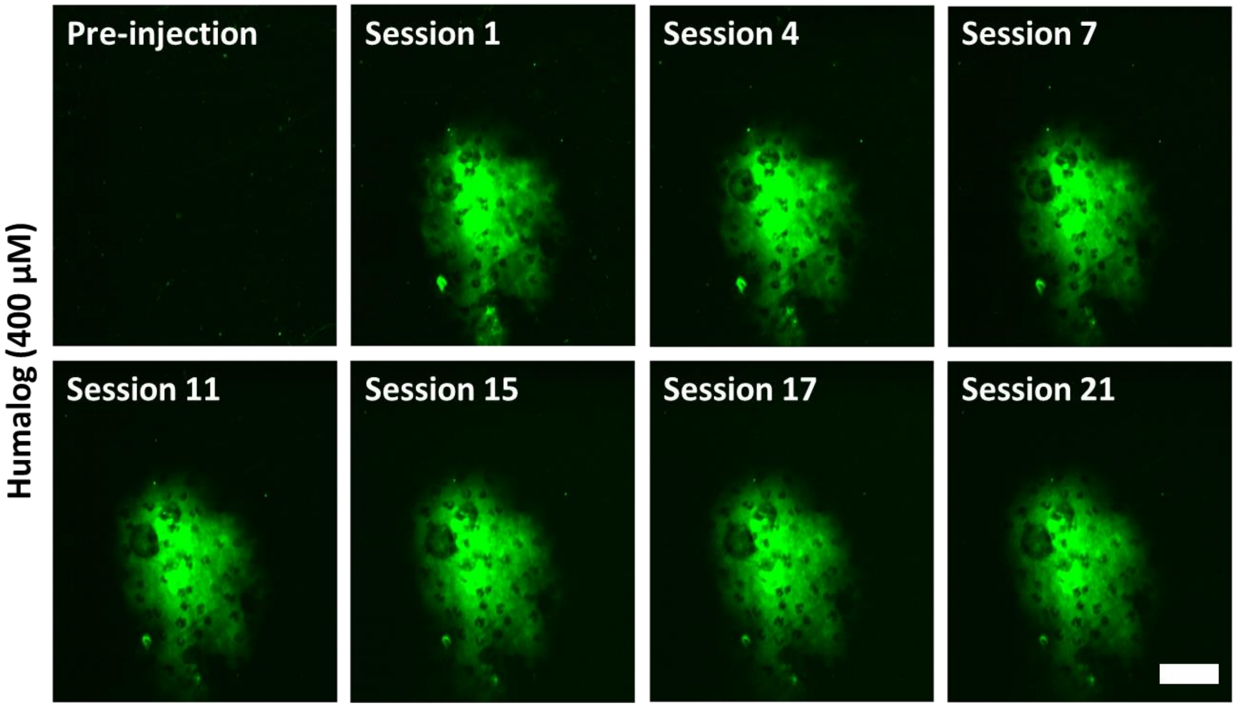


**Figure S1:** Photobleaching test of Cy7.5 dye-labeled lispro (400 µM) in Humalog formulation injected in a dead mouse ear. Over the course of 3 hours, 21 imaging sessions were performed and no decrease of photoacoustic signals were found. Scale bar = 500 µm.

a b

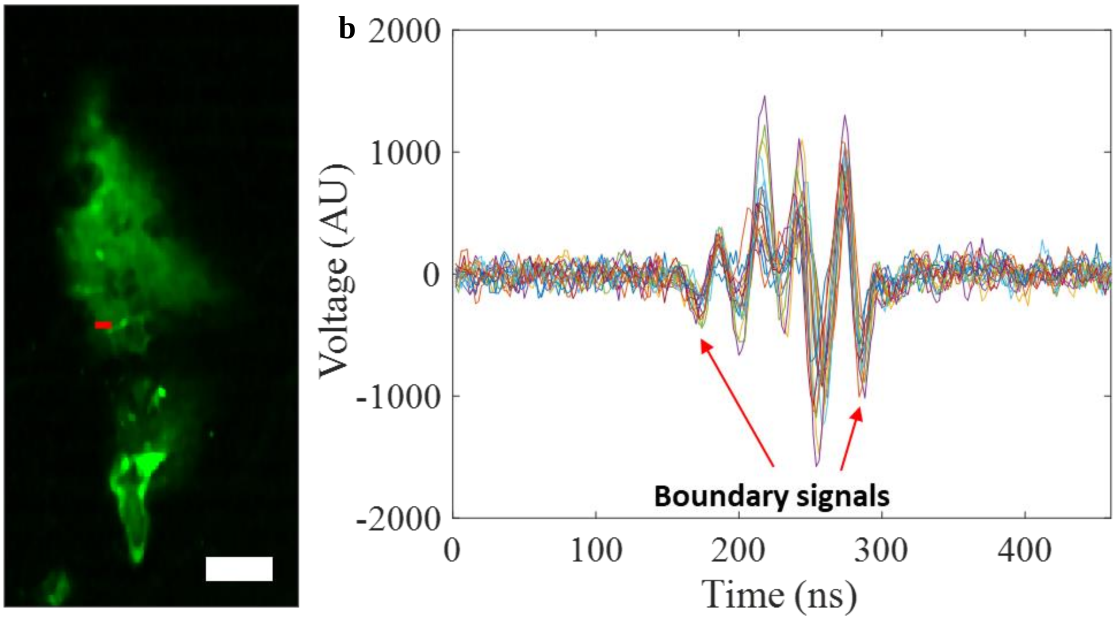


**Figure S2:** (a) Dye-labeled insulin lispro in Humalog formulation injected in the mouse ear. (b) The photoacoustic signals along the red line in image a. The photoacoustic signals within the boundary signals were found to be developed. Due to the limited bandwidth of ultrasonic transducers, the photoacoustic signal from homogeneous samples such as blood in large vessels is averaged out from inside the light-absorbing volume, leading to the detection of only boundary signals [1,2]. For such samples, quantification is usually performed by estimating the total volume from the boundary and determining the concentration as the direct measure of the boundary signal amplitude. However, exogenously injected dye-labeled lispro tends to produce a diffused boundary in biological tissues; therefore, for accurate quantification of the signal, a complete 3D volume must be detected. It appears that the heterogeneous structure of mouse ear tissue provides an inhomogeneous distribution of the dye-labeled lispro at the scale of the central acoustic wavelength of our transducer, producing a well-developed speckle pattern with little boundary buildup [3,4]. Scale bar = 200 µm.

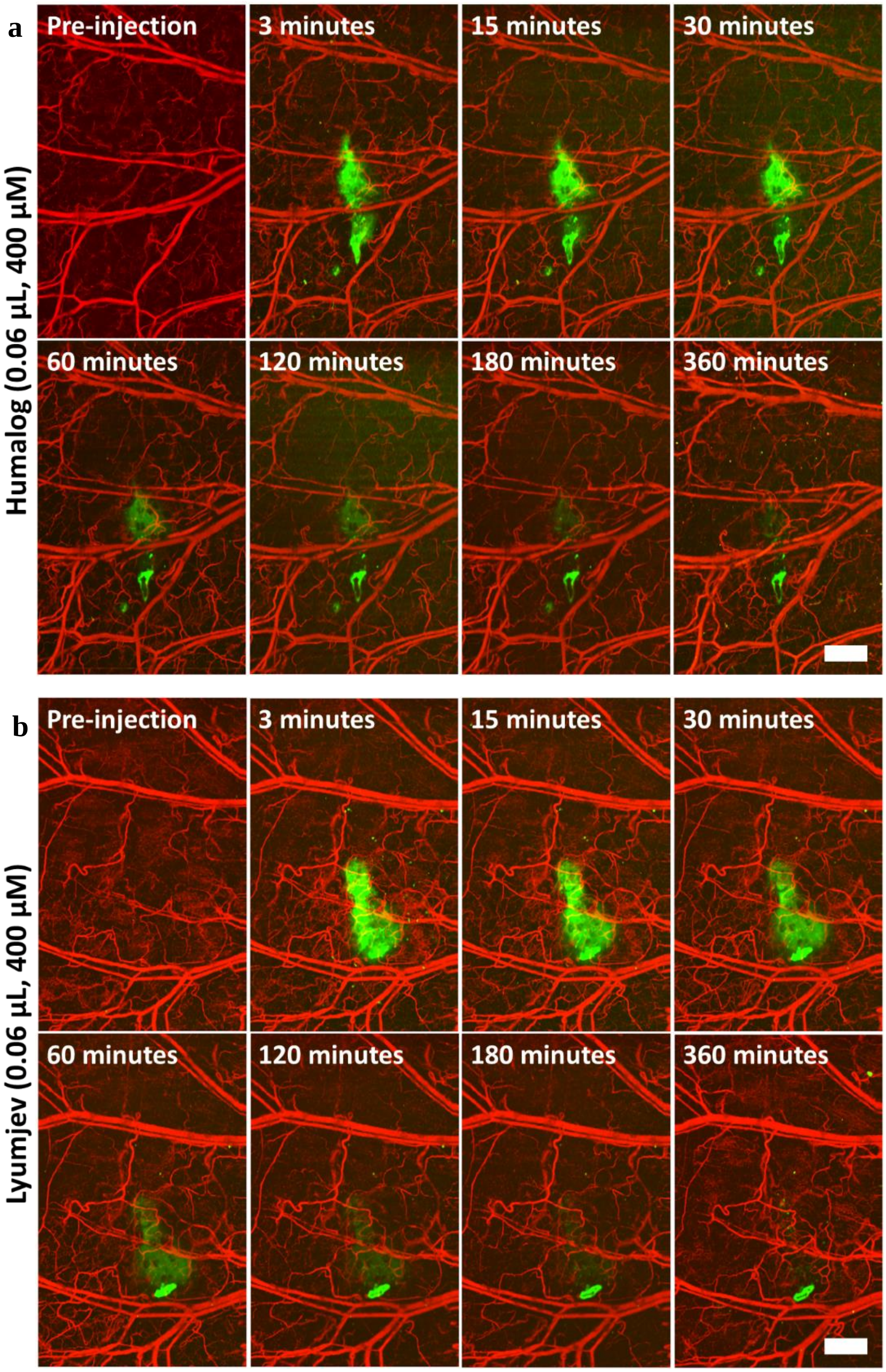


**Figure S3:** Visualization of dynamics of Cy7.5 dye-labeled insulin lispro in the Humalog (a) and Lyumjev (b) formulations (0.06 μL at 400 μM) in the mouse ear. The mouse was kept awake in moving condition between 180 minutes and 360 minutes. Scale bars = 500 μm.

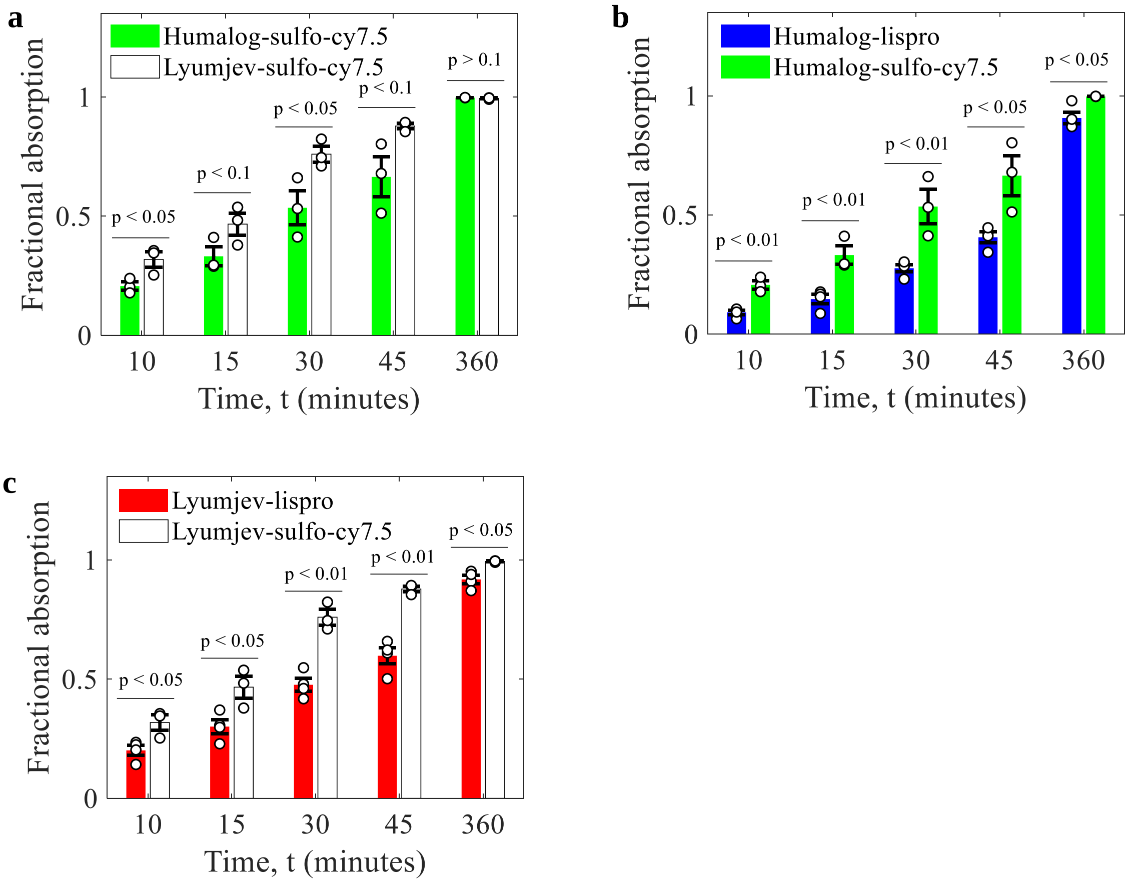


**Figure S4:** (a) Comparison of absorptions of sulf-cy7.5 dye in the Humalog and Lyumjev formulations (n = 3). (b) Comparison of absorptions of dye-labeled insulin lispro (n = 4) and sulfo-cy7.5 dye (n = 3) in the Humalog formulation. (c) Comparison of absorptions of dye-labeled insulin lispro (n = 4) and sulfo-cy7.5 dye (n = 3) in the Lyumjev formulation. All the p-values were calculated by unpaired two-sample t-test after performing Shapiro-Wilk normality test. For comparison of the absorption of dye-labeled insulin lispro with that of sulfo-cy7.5 dye, an F-test was performed to confirm equal variances. In case of unequal variances (only at 360 minutes for both plot b and plot c), an unpaired two-sample t-test considering unequal variances (also known as Welch's t-test) was performed. All data represent mean ± standard error of the mean.

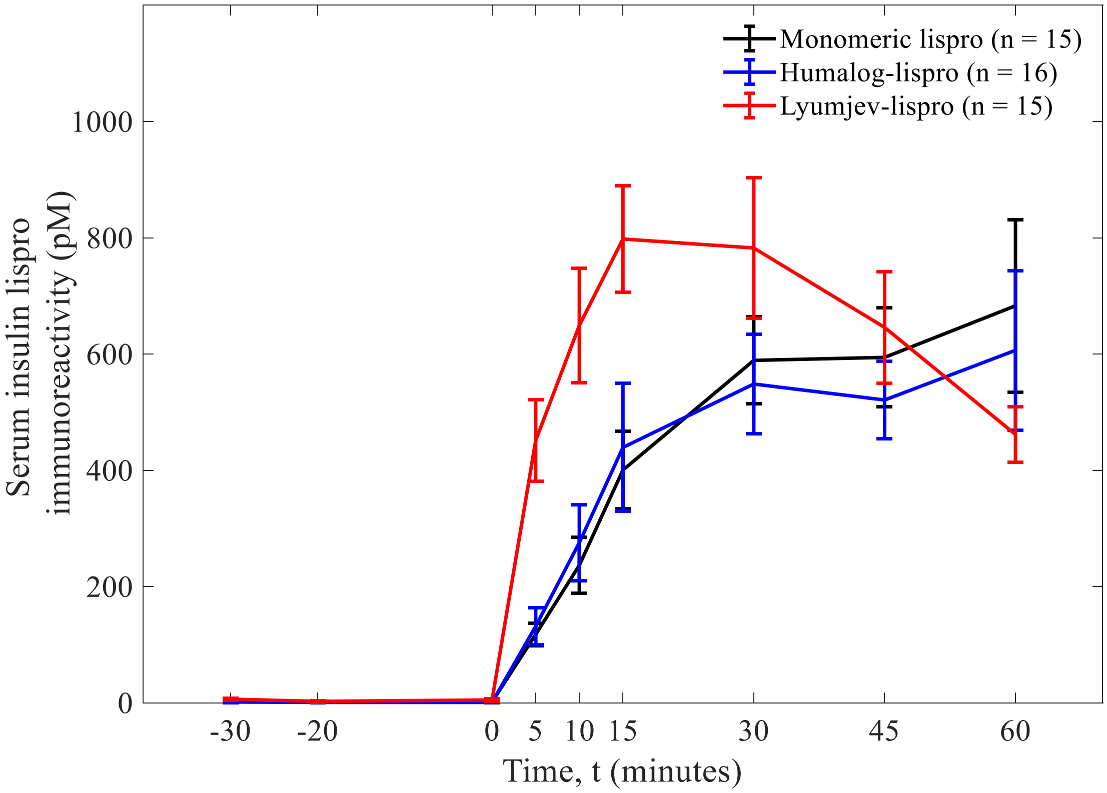


**Figure S5:** Pharmacokinetic profiles of monomeric lispro (n = 15) in Humalog formulation without zinc and phenolic ligand, hexameric lispro (n = 16) in Humalog formulation, and hexameric lispro (n = 15) in Lyumjev formulation in pigs, for the first 60 min post-injection. Data represent mean ± standard error of the mean

**Table S1: Pharmacokinetic parameters (± SEM) of lispro formulations in pigs.**

| Drug | $T_{max}$ (min) | Early $_{1/2\ Max}$ (min) | $C_{max}$ (nM) | $AUC_{INF}$ (nM*min) | $AUC_{0\text{-}30}$ (nM*min) |
|---|---|---|---|---|---|
| Humalog (n =16) | 59 ± 8 | 20 ± 3 | 1.11 ± 0.38 | 104 ± 23.3 | 10.5 ± 1.8 |
| Monomeric lispro (n = 15) | 70 ± 11 | 29 ± 8 | 1.03 ± 0.18 | 104 ± 14.0 | 10.2 ± 1.4 |
| Lyumjev (n = 15) | 22 ± 3 | 6.9 ± 1.0 | 1.01 ± 0.10 | 87.8 ± 7.2 | 19.2 ± 2.2 |

$T_{max}$ = time at maximum observed concentration.

Early $_{1/2\ max}$ = first time the pharmacokinetic curve rises above ½ of $C_{max}$ (maximal observed concentration) value.

$C_{max}$ = maximal observed concentration.

$AUC_{INF}$ = area under the curve from time 0 to infinity.

$AUC_{0\text{-}30}$ = area under the curve from time 0 to 30 min.

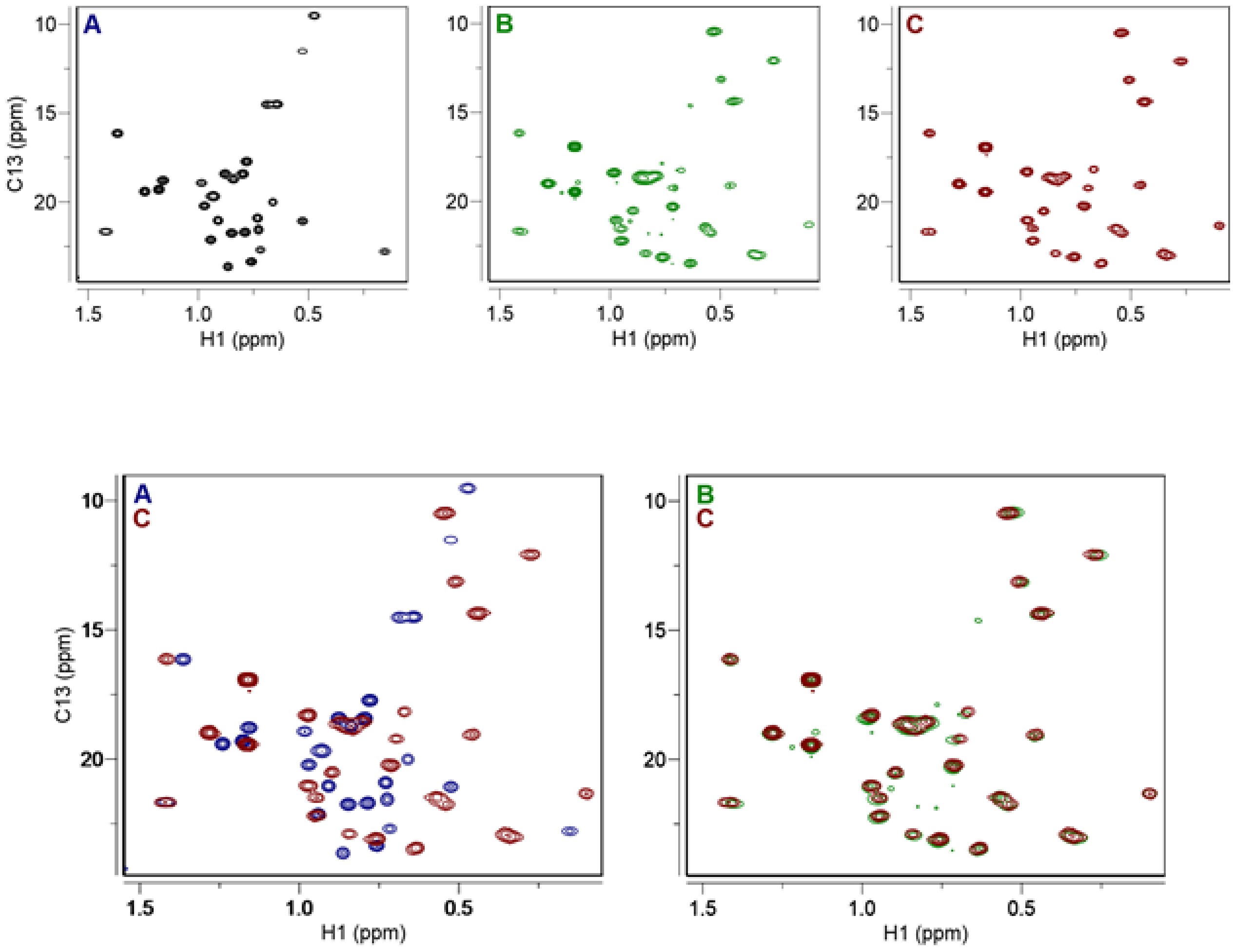


**Figure S6:** (Top) 2D $^{1}H$-$^{13}C$ HSQC NMR spectra expanded to the methyl region of (A) Zinc-free lispro, (B) Humulog U100, and (C) Lyumjev. (Bottom) Overlays of the same spectra.

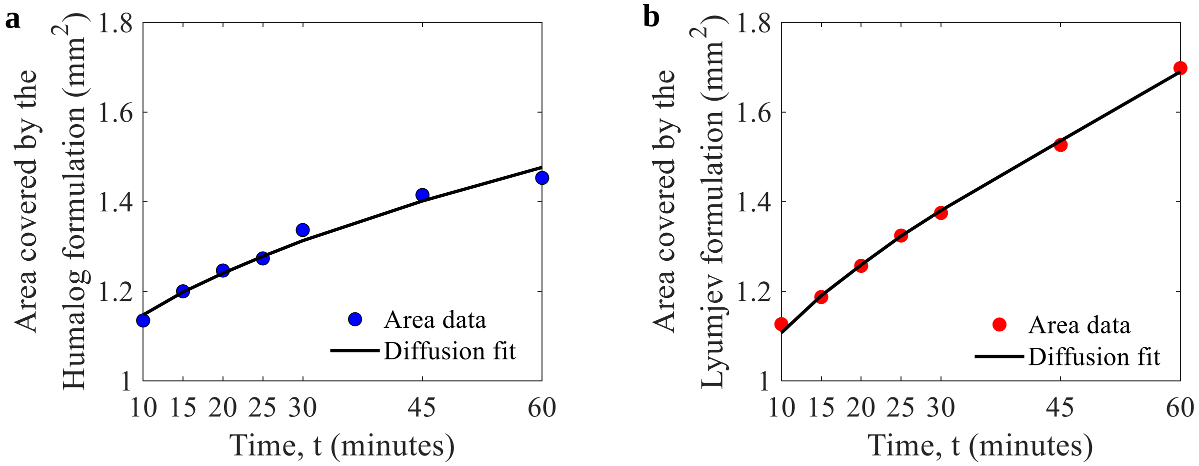


**Figure S7:** (a) Fitting of diffusion curve on the temporal area change data for dye-labeled insulin lispro in the Humalog formulation. (b) Fitting of diffusion curve on the temporal area change data for dye-labeled insulin lispro in the Lyumjev formulation.
